# Enabling spectrally resolved single-molecule localization microscopy at high emitter densities

**DOI:** 10.1101/2022.06.29.498127

**Authors:** Koen J.A. Martens, Martijn Gobes, Emmanouil Archontakis, Roger R. Brillas, Niels Zijlstra, Lorenzo Albertazzi, Johannes Hohlbein

## Abstract

Single-molecule localization microscopy (SMLM) is a powerful technique for elucidating structure and dynamics in the life- and material sciences with sub-50 nm spatial resolution. The simultaneous acquisition of spectral information (spectrally resolved SMLM, sSMLM) enables multiplexing using spectrally distinct fluorophores or enable the probing of local chemical environments by using solvachromatic fluorophores such as Nile Red. Until now, the widespread utilisation of sSMLM was hampered by several challenges: an increased complexity of the optical detection pathway, limited software solutions for data analysis, lower accessible emitter densities or smaller field-of-views, and overall compromised spatio-spectral resolution. Here, we present a low-cost implementation of sSMLM that addresses these challenges. Using a blazed, low-dispersion transmission grating positioned close to the image plane here represented by the camera sensor, the +1^st^ diffraction order is minimally elongated compared to the point spread function of the 0^th^ order and can therefore be analysed using common subpixel single-molecule localization algorithms. The distance between both PSFs provides accurate information on the spectral properties of the emitter. The minimal excess width of 1^st^ order PSFs enables a fivefold higher emitter density compared to other sSMLM approaches whilst achieving a spatio-spectral localization accuracy sufficient to discriminate between fluorophores whose peak emission are less than 15 nm apart as demonstrated using dSTORM, DNA-PAINT and smFRET. We provide an ImageJ/Fiji plugin (sSMLMAnalyzer) and suitable Matlab scripts for data analysis. We envision that our approach will find widespread use in super-resolution applications that rely on distinguishing spectrally different fluorophores under low photon conditions.

## Introduction

Super-resolution microscopy, or nanoscopy, has revolutionized the life and material sciences as it allows surpassing the optical diffraction limit by more than an order of magnitude^1–4^. One frequently used implementation is single-molecule localization microscopy (SMLM), in which the stochastic activation of single fluorescent emitters leads to spatially separated point spread functions (PSFs) that are used to determine the position of each emitter with sub-50 nm accuracy. Localisations obtained via (direct) stochastic optical reconstruction microscopy ((d)STORM)^1,4^, point accumulation for imaging in nanoscale topography (PAINT))^5–8^, or photo-activatable localization microscopy (PALM)^3,9,10^ provide access to detailed structural images, or can quantify dynamics and mobilities via single-particle tracking (spt)^10,11^. In this capacity, SMLM has led to breakthroughs^12,13^ in fields such as DNA-protein interactions^14–17^, cell biology^18–20^, and soft matter^8,21–23^.

Improving throughput via multiplexing of different fluorophores in SMLM^5^, enabling microenvironmental characterisation^24,25^, or studying fluorophore-to-fluorophore distance (via single-molecule Förster Resonance Energy Transfer (smFRET))^26,27^, can be accomplished by combining SMLM with the additional spectral characterisation of emitters. Spectral information of single emitters can be acquired and analysed using various implementations that all rely on placing additional components into the optical detection pathway.

The first implementation of spectrally-resolved SMLM uses the ratiometric distinction of spectral emission profiles, and is based on placing one (or more) suitable dichroic mirror(s) in the emission pathway^28–31^. Photons emitted from the sample are separated based on their wavelength, and directed towards two different detection channels. This entails either two separate detectors or using two areas on the same camera chip after using additional lenses or mirrors to guide the beams. Then, the PSFs and their integrated intensities obtained in the two channels are matched, and the intensity ratio of photons is used to discriminate between the emission spectra of different fluorophores. Importantly, this method requires photons to be directed towards each channel implicating that only a defined spectral range around the cut-off wavelength of the dichroic mirror can be accessed.

The second implementation uses point spread function engineering to obtain spectral information on emitters. Here, a spatial light modulator (SLM) or a phase mask (PM) is employed in the Fourier plane of the emission path^32,33^. The introduced phase offset by these elements is depending on the incoming wavelength, which can be exploited to design a pattern so that different PSF shapes are realised when photons of different wavelength arrive at the detector. However, small spectral emission differences in the order of tens of nanometres in the peak emission cannot create sufficiently distinct PSF shapes, hindering discrimination of spectrally close fluorophores. Moreover, the voltage (for SLM) or phase (for PM) has to be specifically tuned for certain emission wavelengths, complicating this method when different fluorophores are used.

In the third implementation, spectral dispersion, a spectrally-dispersive optical element is added, after which the spatial and spectral profiles are guided to different regions on a single camera chip or to completely separate detectors^34–41^. Generally, the spatial profile is then analysed with regular single-molecule localization algorithms^42–45^, while the spectral profile is spread out over tens of pixels and is used to determine the corresponding emission profile. While this implementation allows a large spectral range to be used and allows the discrimination of fluorophores with similar emission spectra, it has various downsides. First, the entire emission pathway needs to be modified to separate the spatial from the spectral channel^37,46^. Second, as the spectral information is widened over tens of pixels, the signal-to-noise ratio obtainable in this channel is compromised, leading to a loss in spectral accuracy^47^. The wide spreading of emission further directly limits the usable density of emitters in the sample, as overlapping spectral profiles cannot be resolved.

Comparing these implementations, it is obvious that there is a need to combine an easy implementation with a broad spectral range and good specificity. Here, we demonstrate spectrally-resolved singlemolecule localization microscopy (sSMLM) using an inexpensive (blazed) transmission grating that can be easily implemented in most microscope configurations. By using a grating with a large line pitch and placing it close to the image plane to minimise the dispersion of the 1^st^ order, we created a low-dispersion sSMLM implementation in which the 0^th^ and 1^st^ diffraction order of every emitter is imaged in a single field of view without the need for any additional optical elements. Our novel implementation is capable of accurately determining spectral properties of single molecules at much higher emitter densities than other spectral-dispersing sSMLM implementations. With our sSMLM approach, we show the technical feasibility of spectral multiplexing and allowing to distinguish between 0% and 15% FRET efficiency in single-molecule FRET experiments.

## Materials and Methods

### Microscopy hardware for SMLM detection

All measurements were performed on a home-build super-resolution microscope fully described elsewhere^17^. Briefly, the 561 nm and 642 nm laser lines of an Omicron Lighthub 6 (Germany) were employed in HiLo (highly inclined and laminated optical sheet) or TIRF (total internal reflection fluorescence) illumination mode via a Nikon 100x 1.49 NA HP/SR objective (Japan). The emission light was guided via the bypass mode of a rescan confocal microscope (RCM, Confocal.nl, The Netherlands) to an Andor Zyla 4.2+ sCMOS camera (United Kingdom) set to 2 x 2 internal pixel binning leading to an 122 nm effective pixel size. Data acquisition was controlled via micromanager^48^. Either a 405/488/561 dichroic mirror and filter set (ZT405/488/561rpc and ZET405/488/561m-TRF, Chroma, Bellows Falls, VT, USA) for the smFRET experiments, or a 405/488/561/642 dichroic mirror and filter set (ZT405/488/561/642rpc and ZET405/488/561/642m-TRF, Chroma) was used. The second set was only used in experiments where the 642 nm laser line was employed.

### Spectral multiplexing with fixated and immunostained cells (dSTORM)

Immobilized Cos7 cells with CF660-immunostained clathrin and CF680-immunostained microtubulin were obtained from Abbelight (France). Cells were imaged for 100.000 frames at 20 ms frame time using 80 mW of laser power at 642 nm excitation wavelength. A STORM buffer (Abbelight, France) was added directly before sealing the sample, which was ~5 minutes before the start of imaging. Data acquisition was started after ~2 minutes of continuous laser illumination used to switch most molecules from an on-state to their off-state thereby lowering the density of emitters.

### Spectral multiplexing with polystyrene nanoparticles (DNA-PAINT)

Streptavidin-coated polystyrene nanoparticles (NPs) with a diameter of 400-700 nm (SVP-05-10; Spherotech, Lake Forest, IL, USA) were loaded with single-stranded DNA (ssDNA) by mixing a biotinylated ssDNA strand (Integrated DNA Technologies, Coralville, IA, USA) with the NPs for 1 hr at room temperature, either with docking strand 1 (NP1) or docking strand 2 (NP2)^49^. Two oven-cleaned (heated to 500 °C for 20 minutes to remove organic impurities) precision #1.5H coverslips (Paul Marienfield GmbH, Germany) are used to create a small ‘chamber’ that is open at two ends, separating the coverslips with double-sided tape. This chamber is filled with a 0.1 mg/mL BSA, 10mM Tris-HCl, 100 mM NaCl, pH 8 solution for 1h, washed with PBS, and filled with PBS containing 0.25 mg/mL NP1, 0.25 mg/mL NP2 for 10 minutes. Afterwards, the chamber is washed with PBS, and 5 nM of each imager strand (complementary to either NP1, containing ATTO655, or complementary to NP2, containing ATTO647N (both Eurofins Genomics, Germany); Supplementary Table 1) in a pH 8 solution containing 5 mM Tris-HCl, 10 mM MgCl2, 1 mM EDTA. The sample is then sealed with tape. Imaging was performed for 2500 frames at 100 ms frame time using a 642 nm laser power of 80 mW in TIRF mode, without any additional emission filter present.

### Spectrally-resolved single-molecule Förster resonance energy transfer (smFRET)

Biotinylated double-stranded DNA labelled with ATTO550/ATTO647N with 23-bp (15% FRET) or 15-bp (55% FRET) separation was obtained via a multi-lab study on FRET measurements (1-lo, 1-mid)^27^.

PEGylated and biotinylated coverslips were created following an earlier protocol^50^. Briefly, oven-cleaned (heated to 500 °C for 20 minutes to remove organic impurities) coverslips were washed in acetone, and incubated in 1:50 Vectabond/acetone solution (Vector labs, Burlingame, CA, USA). Wells with around 25 μl capacity were created using silicone culture well gaskets (6 mm diameter, Grace Bio-Labs, Bend, OR, USA), and incubated with 200 mg/ml NHS-PEG (Laysan Bio, Arab, AL, USA) and 2.5 mg/ml NHS-biotin-PEG (Laysan Bio) in 50mM MOPS buffer. Then, ~20 μl 0.02 mg/ml neutravidin was added, and after rinsing with 2x 200 μL PBS, 20 pM of the biotinylated DNA was added. The samples were rinsed with 2x 200 μL PBS, and an oxygen scavenger system was added^51,52^ (final concentrations: 1 mM Trolox, 1% glucose oxidase/catalase, 1% glucose). Finally, the gaskets were sealed by placing a coverslip on top. Four movies of 2500 frames each at 100 ms frame time were recorded using ~20 mW laser power (561 nm excitation wavelength) in TIRF mode without an additional emission filter.

### Low dispersion diffraction grating for sSMLM

A blazed transmission grating (70 grooves/mm, Edmund Optics, Barrington, NJ, USA, part nr 46-067) was housed in a 3D-printed plastic insert (Supplementary information) and was inserted in the inner tube of the camera. An external C-mount threaded retainer ring (Thorlabs, Newton, NJ, USA, part id CMRR) was then threaded to retain the grating in this place, and the retainer ring was glued to the plastic insert for repeatable insertion of the grating. The grating was inserted such that all diffractions orders from a single emitter are on the same horizontal level. In our implementation, the +1^st^ representing the spectral channel was to the right of the 0^th^ order. Angular deviations from the expected tilt of 0 radians can be accounted for by the software discussed below. As given by the manufacturer, the transmission grating has an overall efficiency of 73% at 632.8 nm wavelength with 41% and 32% for the 0^th^ and +1^st^ order, respectively.

While the absolute distance between the camera chip and the grating is initially unknown, a full rotation of the C-mount thread corresponds to exactly 1/32^th^ inch (~0.8 mm). Four sSMLM experiments with increasingly more full rotations of the grating away from the camera were recorded. The median value of the resulting distance between the 0^th^ and the 1^st^ order is plotted against the absolute distance that the grating has been moved from the closest position (Supplementary Figure 2). Fitting and extrapolating this curve with a 1^st^ order polynomial reveals the distance of the grating to the chip at the crossing of the curve with the *x*-axis.

### Analysis of sSMLM data

The dSTORM and DNA-PAINT single-molecule datasets were localized via ThunderSTORM^53^ for Fiji^54,55^, with the pSMLM plugin^44^, after performing a 50-frame temporal median filter^56^. A β-spline wavelet filter with scale 2 and order 3, a local maxima finder with a threshold set to the standard deviation of wave F1 of the filter multiplied by 1.5, and a non-calibrated 3D astigmatism Gaussian fitting routine (dSTORM) or a pSMLM routine with ROI (region of interest) 11-x-11 pixels (DNA-PAINT), respectively, was used. For analyses where the width of the PSFs was critical (i.e. the smFRET experiments), analysis was performed by SMAP with the fit3dSpline fitter^43,57^ without a calibrated PSF model. In SMAP, a difference-of-Gaussian filter with size 3 pixels was used alongside an absolute photon cut-off value of 0.3 with a 5-pixel NMS kernel size, to identify PSFs. Fitting was performed with an elliptical PSF with a 13-pixel ROI, 30 iterations. No further filtering was performed.

As the grating leads to PSF pairs representing the 0^th^ order and +1^st^ diffraction order, an algorithm (in MATLAB 2019b (The MathWorks, Natick, MA, USA) or JAVA) needs to identify which pairs belong to an individual emitter. We used a similar analysis as used for linking and analysing individual lobes of a double-helix point spread function^58^. First, possible pairs were found for a given distance and rotation regime from all localizations on a single frame. Rotation and distance bounds can be determined either by hand from the raw images or algorithmically (Supplementary Note 1). Next, localizations that only have a single possible pairing partner were collapsed and refrained from further linking. This is repeated until all pairs are collapsed, or until no further pairs can be collapsed. In the latter case, non-paired localizations were removed from further analysis. Finally, the position of the spatial (here, left-most) localization, the distance and angle between the two localizations, and (if applicable) the obtained PSF width in both dimensions for the spatial and the spectral localizations were stored. The code belonging to this algorithm is provided as supplemental data.

### Analysis of spectrally resolved smFRET data

For the spectral smFRET analysis, we used the workflow as presented above while using SMAP with the fit3dSpline fitter for the sub-pixel localisation determination. In addition to the 0^th^ to 1^st^ distances, we obtained the spectral widths of the 1^st^ order of all pairs and plotted this data as a 2D histogram (i.e., spectral distance versus spectral width). The histograms of both 15% and 55% FRET measurements were combined, and fitted with 4 Gaussian profiles, representing donor-only, 15% FRET, 55% FRET, and background populations (not shown) corresponding to spurious or noisy localizations. These Gaussian profiles were re-fitted to either the 15% FRET or the 55% FRET 2D histograms, only changing the relative intensities of the profiles, but not their position or width.

Thereafter, individual linked pairs (i.e., a 0^th^ and 1^st^ order pattern) were classified as ‘FRET’, ‘Donor Only’, or ‘Background’, based on the likelihood of belonging to each Gaussian profile. Individual linked pairs were furthermore linked to other linked pairs throughout time via a simple Nearest-Neighbours tracking algorithm^59^, with maximum 1 pixel (122 nm) movement between frames. Only tracks with at least 10 localisations without donor blinking or bleaching events were investigated further.

### Simulating sSMLM emission data

Simulation of the expected spatial and spectral diffraction patterns was performed in MATLAB 2019b based on the physical properties of the grating and its placement in respect to the camera. First, the emission spectrum on which the simulation is based is quantised in single wavelength units (i.e., a resolution of 1 nm). For every wavelength, the angle between the 0^th^ and the +1^st^ order diffraction towards the grating is calculated based on the specified density of grooves of the grating (here 70 mm^-1^). The angle is then used to obtain the spatial position of the 1^st^ order diffraction pattern on the camera chip (based on a specified distance of the grating to the detector). Then, a PSF is approximated via a 2-dimensional Gaussian function, positioned at the 0^th^ order and the 1^st^ order diffraction pattern positions, where the positions are offset by a pre-defined random position between 0 and 1 final pixel size to accurately account for the random positions of emitters in SMLM. The simulated profiles are then normalized with respect to their relative intensity (specified by the emission profile and by the efficiencies of the diffraction orders (41% and 32% for the 0^th^ and 1^st^ order, respectively)). This simulation is first over-sampled on a grid with 1 nm^2^-sized pixels, before it is binned into 122 nm by 122 nm pixels, representing condition of our sSMLM hardware. The code belonging to this simulation is provided as supplemental data.

### Simulating the resolvable emitter density

The resolvable emitter density possible in our implementation was simulated in MATLAB 2019b with similar conditions as performed previously^37^. A 20-by-20 μm frame was filled with emitters specified by a certain density. Then, localizations that are located closer apart than 3 pixels (here 0.366 μm) are indicated as ‘overlapping’. For our sSMLM implementation, a secondary localization was placed to the right of the primary emitter with a randomly chosen distance between 2500 and 3400 nm, and was taken into consideration for overlapping scenarios. This was repeated 500 times at every density to determine the mean and the standard deviation of each condition.

### Spectral dispersion characterisation

A 2D DNA-PAINT GATTA-PAINT 80RG (Gattaquant, Germany) sample containing ATTO542 and ATTO655 fluorophores attached to imager strands was imaged with either a 561 nm or a 642 nm laser activated. The 0^th^-to-1^st^-order distance was calculated, and the difference in the median distance was divided by the difference of the weighted mean of the fluorophore emission profiles. The fluorophore emission profiles were corrected for the optics and detector (specifically: for the dichroic mirror and filter set, emission filter, and sCMOS quantum efficiency) used in the microscopy system^17^.

## Results

### Implementation and characterisation of minimal dispersion sSMLM

To maximise the signal-to-noise ratio and the achievable molecular density in spectrally-resolved single-molecule localization microscopy (sSMLM), the available photon budget should be distributed over as few camera pixels as possible^47^. To fulfil this criterium, we placed a transmission grating with low dispersion (70 lines/mm) as close as possible to a camera chip representing the image plane (< 1 cm, Fig. 1a) by placing it inside the camera housing. This arrangement minimises the separation of the 0^th^ and 1^st^ order diffraction patterns, and thus results in the highest achievable fluorophore density and signal-to-noise ratio for the 1^st^ order diffraction pattern (Fig. 1b, Fig. 1c). Notably, and distinctly different from earlier implementations^34–37^, our arrangement allows imaging of both spatial and spectral information in the same field of view, thereby maximising the usable detection area on the sensor. Furthermore, our approach does not require any additional optical components such as mirrors, beam splitters, or secondary detectors.

**Figure 1:**
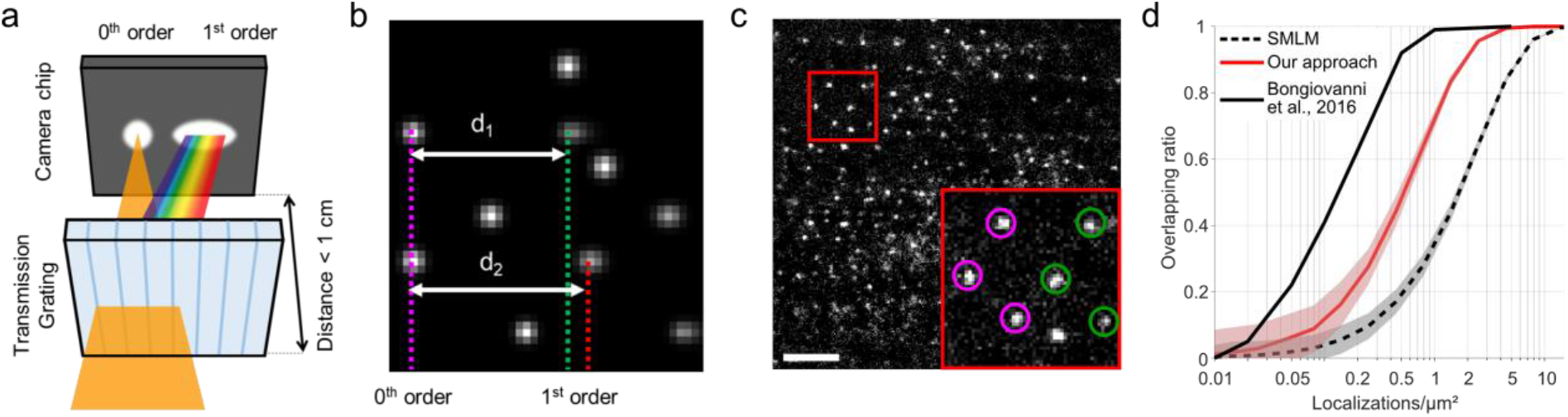
Implementation of low-dispersion spectrally-resolved single-molecule localization microscopy. **a)** A low-dispersion blazed transmission grating is placed in the emission path of a typical SMLM capable microscope such that the distance of the grating to the image plane is minimised. Around 50% of the light passing the grating will not have any dispersion, causing a 0^th^ order point spread function (PSF) to appear. The other 50% of the light is dispersed based on its wavelength and will create a second, slightly elongated 1^st^ order diffraction pattern. Image not to scale. **b)** Simulation of low-dispersion sSMLM data of two spectrally different emitters with λ_1_ (corresponding to d_1_) < λ_2_ (corresponding to d_2_). Six emitters create in total 5 0^th^ and 1^st^ order diffraction pairs on the detector, which can be linked together (two 1^st^ order diffraction patterns are not captured in the field of view). The obtained distances between the 0^th^ and 1^st^ order diffraction patterns (d_1_ and d_2_) are a measure for the average emission wavelength. **c)** Single frame of raw data of a DNA-PAINT nanoruler sample showing 85 spatio-spectrally resolvable emitters in a 31 x 31 μm field of view. The red outline is enlarged in the inset, in which the 0^th^ and 1^st^ order diffraction patterns are encircled in magenta and green, respectively. Scalebar represents 5 μm. **d)** Comparison of achievable emitter density in standard (non-spectrally resolved) SMLM (black dotted line), our approach (red line), and sSMLM with 20-30 pixels spectral pattern elongation, taken from Bongiovanni et al.^37^. The shaded background indicates the standard deviation as determined from repeating the simulations.

As the PSFs from the obtained 1^st^ order diffraction pattern show only minor excess width compared to the PSFs in the 0^th^ order, we were able to employ existing super-resolution algorithms to independently obtain sub-pixel localizations of the 0^th^ and 1^st^ order diffraction patterns. Next, we linked the localizations with each other in the dispersion direction, with the distance between the 0^th^ and 1^st^ order diffraction patterns (*d*1 and *d*2 in Fig 1b; further called ‘0^th^-to-1^st^-order distance’) being a direct measure for the average emission wavelength of the emitter (*λ*_1_ < *λ*_2_ with *d*_1_ < *d*_2_). Moreover, the excess width of the 1^st^ order diffraction pattern compared to the width of the PSF in the 0^th^ order is a measure for the width of the emission spectrum. This directly results in a spectral accuracy being limited primarily by spatial single-molecule localization accuracy, an area of research which is progressing very rapidly via both software and hardware developments^42,60,61^. Using minimal dispersion, the achievable density in our implementation without the need for specialized high-density fitting algorithms is around 5 times higher than earlier implementations of sSMLM, where the spectral information is spread out over 20-30 pixels (Fig. 1d, Methods)^37^.

We determined the distance between the dispersion-inducing optics of the grating to the camera chip to be 6.9 ± 0.1 mm in our system (Methods, Supplementary Fig 1a,b). The spectral dispersion (SD) was determined by calculating the 0^th^-to-1^st^-order distance of a sample labelled with ATTO542 and ATTO655 (Supplementary Fig 1c). From the median distance of these obtained distances and the mean emission profile of the fluorophores, a spectral dispersion of ~0.21 nm/nm (spatial/spectral; equivalent here to ~27 nm/px (spectral/spatial)) was determined. We did not observe a wavelength dependency on the angle between the spatial and spectral profiles (Supplementary Fig 1d).

Multiplexing of 10 nm spectrally separated fluorophores in dSTORM and DNA-PAINT The signal-to-noise ratio of the obtained 1^st^ order diffraction pattern, combined with high resolution of sub-pixel localization algorithms indicate that small spectral differences can be elucidated. We imaged double-labelled fixated Cos7 cells, in which clathrin was labelled with the fluorophore CF660, and tubulin with CF680. Pseudo-colour coding a super-resolved image based on 0^th^-to-1^st^-order distance reveals good separation of the labelled structures without further analysis (Fig. 2a), even though these fluorophores only have a ~10 nm intensity-weighted spectral separation in our microscope (CF660: 691.9 nm, CF680: 701.6 nm, Fig. 2b; 12 nm difference in peak wavelength (CF660: 686 nm, CF680: 698 nm)).

**Figure 2:**
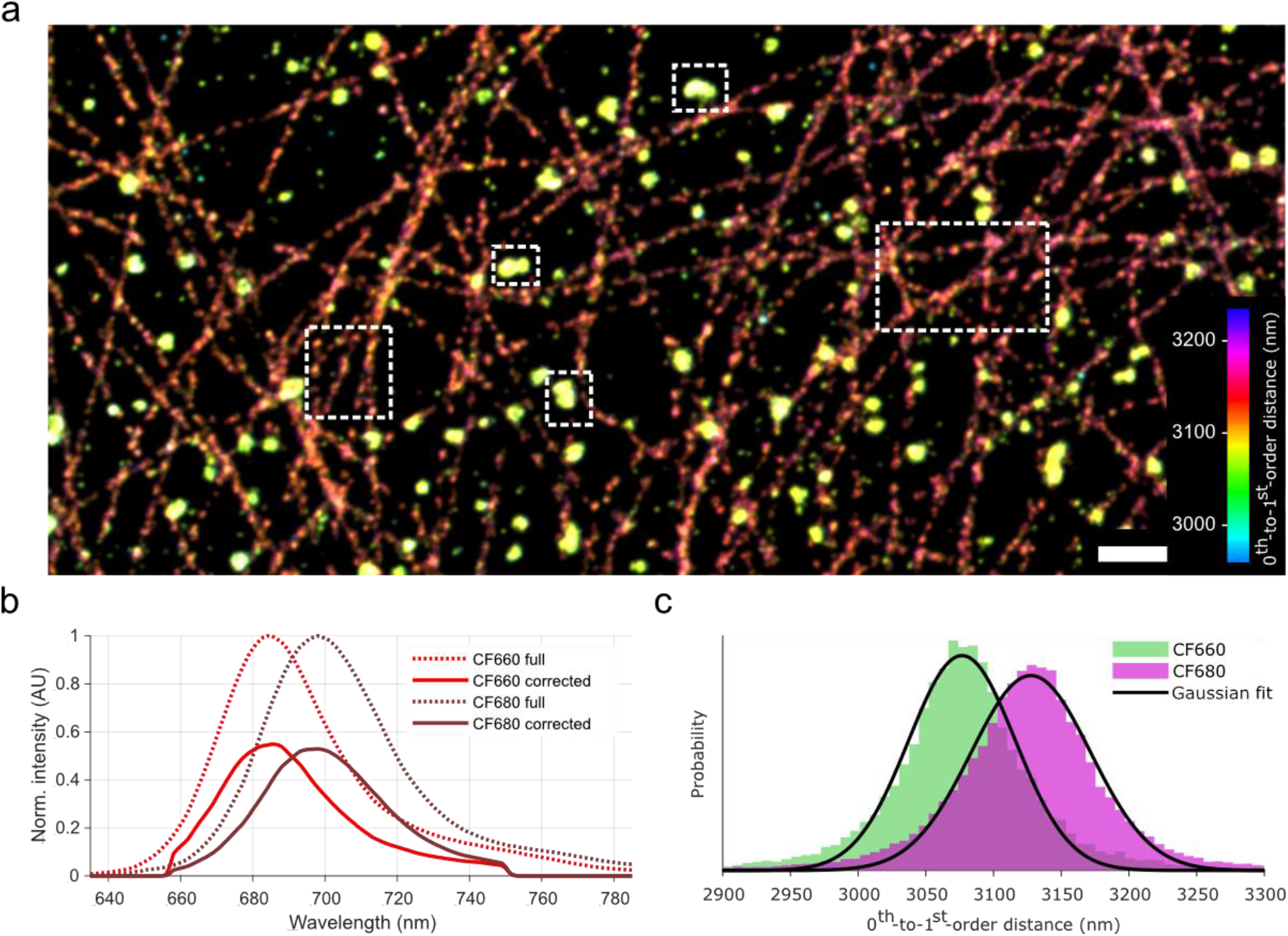
Multiplexing dSTORM of fixated Cos7 cells with CF660-labelled clathrin and CF680-labelled tubules. **a)** Obtained dSTORM image, colour-coded on 0^th^-to-1^st^-order distance. Separation between tubule and clathrin can be observed without further data analysis. Scalebar represents 1 μm. **b)** Emission spectra of CF660 (bright red) and CF680 (dark red). Dotted lines represent full spectra, while the solid lines represent emission spectra corrected for the transmission characteristics of the optical components in the detection path of the microscope. **c)** Histograms representing 0^th^-to-1^st^-order distances of fluorophores belonging to areas indicated by dotted outlines in a. These populations are fitted with Gaussian curves (see main text), and attributed to CF660 (green) or to CF680 (magenta).

Selecting image regions with mostly CF660- or CF680-labelled structures (dotted outlines in Fig. 2a), and fitting the corresponding 0^th^-to-1^st^-order distances with a Gaussian profile reveals that CF660 has a 0^th^-to-1^st^-order distance of 3077 ± 2 nm (σ = 56 ± 2 nm; mean ± 95% confidence interval (CI)), and 3128 ± 2 nm for CF680 (σ = 62 ± 2 nm; Fig. 2c). This is a difference of 51 ± 2 nm in the raw data, which corresponds to a spectral distance of 10.9 ± 0.4 nm, in agreement with the weighted average and peak position difference of the microscope-corrected emission profiles.

Next, we performed a DNA-PAINT experiment on polystyrene nanoparticles (NPs) that have DNA-PAINT docking strands for either ATTO647N or for ATTO655 (Fig 3a). These fluorophores have a weighted average emission wavelength separated by only ~9 nm (684.5 and 693.4 nm, respectively, after correcting for optical components in our microscope (Fig 3b)), and a peak emission wavelength separation of ~14 nm (665 and 679 nm, respectively, after correcting for optical components in our microscope (Fig 3b)). After isolating localizations belonging to individual beads and analysing the 0^th^-to-1^st^-order distances of these emitters, two populations can be observed (Fig. 3c).

**Figure 3:**
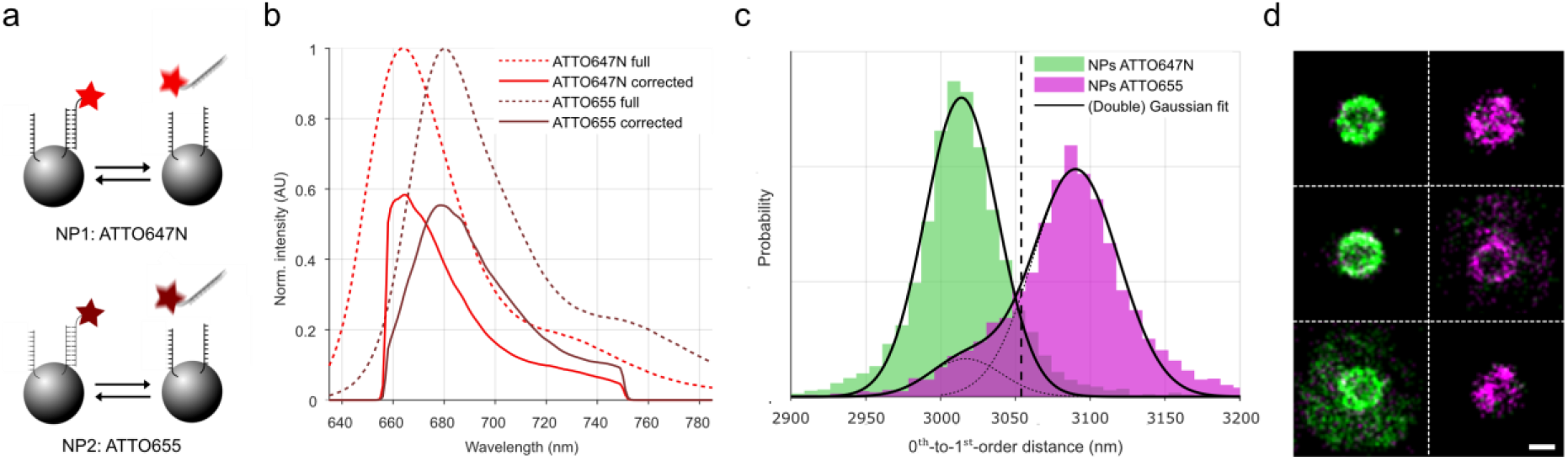
Low-dispersion sSMLM is capable of distinguishing ATTO647N from ATTO655 in DNA-PAINT. **a)** Two different nanoparticles have associated DNA-PAINT imager strands containing either ATTO647N or ATTO655. Scheme not to scale. **b)** Emission spectra of ATTO647N (bright red) and ATTO655 (dark red). Dotted lines represent full spectra, while the solid lines represent emission spectra corrected for the optical components present in the microscope. **c)** Histograms representing observed 0^th^-to-1^st^-order distances of fluorophores belonging to individual NPs. These populations are fitted with Gaussian curve(s) (see main text) and attributed to NPs accepting ATTO647N-DNA (green) or to NPs accepting ATTO655-DNA (magenta). **d**) Visualisation of six individual NPs, with individual localizations colour-coded based on the dotted line shown in b. Scale bar represents 500 nm.

The population with the lowest 0^th^-to-1^st^-order distance (green; Gaussian fit peak position: 3014 ± 1 nm, σ = 34 ± 1 nm, mean ± 95% CI,) was attributed to ATTO647N fluorophores. The population with the larger distances (magenta) was fitted with a combination of the Gaussian curve fitted to the first observed population, along with a unique Gaussian curve (Gaussian fit peak position: 3090 ± 2 nm, σ = 41 ± 3 nm). This population was attributed to ATTO655 fluorophores. The larger standard deviation of the ATTO655 population compared to the ATTO647N population can be attributed to a lower median localization accuracy (42 nm vs 50 nm), possibly caused by a difference in quantum yield (65% vs 30%). The spectral distance between these fitted peak positions (76 ± 2 nm distance; corresponding to 16.2 ± 0.4 nm spectral separation) is close to the difference between the emission peaks of both fluorophores, but higher than the weighted average wavelength. This is possibly caused by deviations of the described wavelength-dependant efficiency of optical elements compared to our hardware implementation, which could lead to a shifted weighted mean emission wavelength of ATTO647N, as its emission maximum is close to the lower spectral cut-off (~660 nm) of the filters and beam splitters used.

Next, all linked localizations were colour-coded according to their distance (cut-off at the black dotted line in Fig. 3c at 3054 nm). Visualisation of the individual NPs (Fig. 3d) then reveals their fluorophore distribution. This shows that the NPs are populated by either one DNA-PAINT docking strand or the other, with minimal cross-talk between the used fluorophores, which can be attributed to unspecific DNA-DNA interactions.

Taken together with the dSTORM data of the fixated cells, the obtained order of the mean emission wavelength of these four fluorophores (ATTO647N, CF660, ATTO655, CF680) coincided with that of the 0^th^-to-1^st^-order distance (calculated mean emission wavelength from spectra provided by the manufacturers and corrected for the optical properties of our setup): 685, 692, 693, 701 nm; mean 0^th^-to- 1^st^-order distance: 3014, 3077, 3090, 3128 nm). The separation of these 0^th^-to-1^st^-order distances suggests that simultaneous multiplexing of at least three fluorophores with single-wavelength excitation is technically possible under realistic experimental conditions (Supplementary Figure 2).

### Elucidation of single molecule FRET states via sSMLM

Next, we were interested whether we could expand the low-dispersion sSMLM to assess single-molecule Förster resonance energy transfer (smFRET). In a typical surface-based smFRET experiment, probes labelled with a donor and an acceptor fluorophore are immobilized and monitored over time. Depending on the experiment, changes in FRET and/or changes in acceptor/donor activity (i.e., blinking and bleaching) can be expected. Conventionally, a ratiometric spectral determination method is applied to separate the donor emission from the acceptor emission on different positions on a camera chip, and the intensity ratio between these is a measure for the relative FRET efficiency *E*. In our implementation, we can use the full field of view of our camera and determine *E* via the 0^th^-to-1^st^ order distance. We further can utilise the width of the 1^st^ order diffraction pattern as an additional way of discriminating FRET.

We performed smFRET measurements on well-characterised samples of immobilized double-stranded DNA that is dual-labelled with ATTO550 and ATTO647N^27^. Two samples with different distances between the labelling sites were used: 23-bp separation (~8.4 nm) and 15-bp separation (~5.9 nm), leading to FRET efficiencies of ~0.15 and ~0.55, respectively. First, we performed simulations using the known emission profiles corrected for the fluorophore’s quantum yield and the optical elements in our microscope (Methods; Fig. 4a,b). These simulations of donor-only, 15% FRET, and 55% FRET samples show that the 0^th^-to-1^st^-order distance follows *d*_donor_ < *d*_15%_ < *d*_55%_, and the width of the 1^st^ order diffraction pattern follows *σ*_donor_ < *σ*_15%_ < *σ*_55%_.

**Figure 4:**
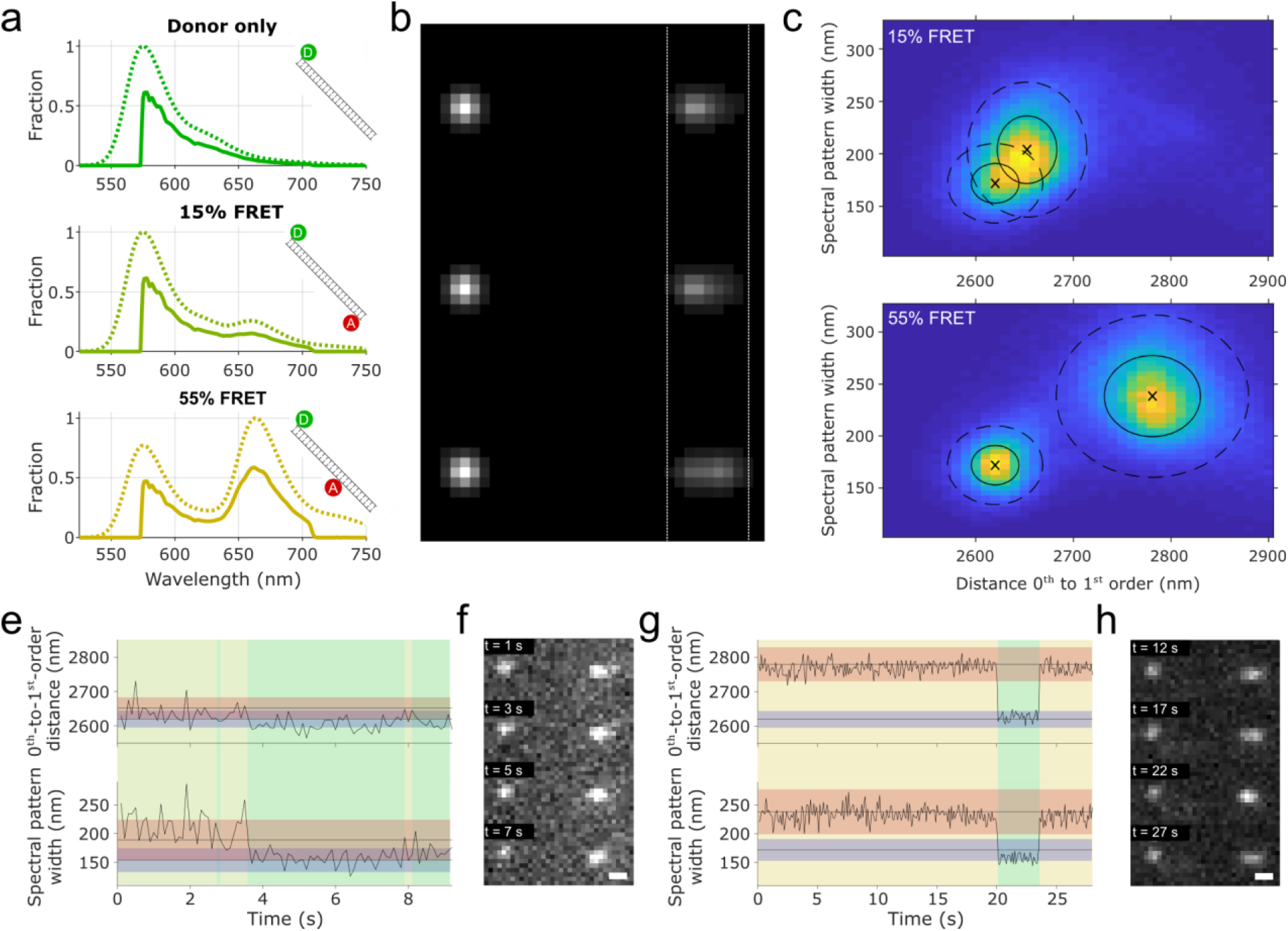
Single-molecule FRET analysis with low-dispersion sSMLM. **a)** Predicted emission spectra of donor-only (top, ATTO550), 15% FRET (middle), and 55% FRET (bottom). Dotted lines represent full spectra, while the solid lines represent emission spectra corrected for the transmission characteristics of the optical components present in the microscope. Schemes represent donor (green; ATTO550) and acceptor (red; ATTO647N) fluorophore placements on a DNA strand. **b)** Simulated raw data obtained in our low-dispersion sSMLM implementation, based on the emission profiles determined in a. Vertical dotted white lines are a guide for the eye. **c,d)** 2-dimensional histograms of experimental data of 15% FRET (c) and 55% FRET (d). The histograms were globally fitted with multiple Gaussian functions (shown here centred around black crosses, with solid ellipses representing 1 sigma, and dotted ellipses representing 2 sigma; Methods, main text). **e-h)** Single emitter time trace analysis of a bleaching acceptor fluorophore in a 15% FRET pair (e,f), and of a blinking acceptor fluorophore in a 55% FRET pair (g,h). Horizontal grey lines with red and blue shading represent 15% (e) or 55% (g) FRET populations (red) and donor-only populations (blue), determined from the fit in c,d. The vertical green, yellow, orange shading represented current FRET pair state, with green representing donor-only, yellow representing 15% FRET, and orange representing 55% FRET. The raw data corresponding to these FRET pairs throughout the observation time is shown in f,h. Scalebars in f,h represent 500 nm.

Experimentally, the immobilized DNA strands were imaged separately. Contrary to the multiplexing before, both the 0^th^-to-1^st^-order distance and the width of 1^st^ order diffraction pattern are measures for the FRET efficiency and were therefore visualised (Fig. 4c,d). The experimental data agrees with the simulations showing *d*_donor_ < *d*_15%_ < *d*_55%_ (2620 nm, 2653 nm, 2781 nm, respectively) and *σ*_donor_ < *σ*_15%_ < *σ*_55%_ (172 nm, 204 nm, 238 nm, respectively).

Next, we explored to which extend we can monitor dynamic behaviour using spectrally resolved smFRET. While no direct state transitions are expected for this sample, there is occasional acceptor fluorophore blinking or bleaching, leading to a transition of FRET emission to donor-only emission. For this, we fitted the combined [d, σ] 2-dimensional histogram with four 2-dimensional Gaussian profiles (Fig. 4c,d black crosses and ellipses). These profiles comprise donor-only, 15% FRET efficiency, 55% FRET efficiency, and ‘background’ states (background state not shown). The ‘background’ state is attributed to nonsense linkages occurring from sparse localizations unrelated to the FRET sample.

Time traces of individual emitters were further assessed (Fig. 4e-h). The likelihood of an emitter belonging to the pre-determined states was calculated (Methods), and the most likely state dictates the background colour of the graphs in Fig. 4e and 4g. With this methodology, we were able to determine acceptor bleaching (Fig. 4e) and acceptor blinking (Fig. 4g) in 15% and 55% FRET experimental data. Accurate state determination of the 15% FRET measurement proved to be difficult due to the overlapping Gaussian profiles representing the FRET and donor-only states (Fig. 4c), whereas this was better discriminable for 55% FRET.

## Discussion

We have demonstrated minimal-dispersion spectrally-resolved SMLM (sSMLM), which fundamentally maximises signal-to-noise and emitter density due to lowest possible photon spread on the detector. In our implementation, we used a single optical component add-on to the detection path leading to a spectral dispersion of just ~0.2 nm/nm (spatial/spectral), orders of magnitude lower than typical grating-based sSMLM implementations. With this implementation, we realised a five times increased emitter density compared to similar approaches, achieved good separation of emitters with a ~10 nm intensity-weighted spectral difference in STORM, and were able to observe changes between 0%, 15%, and 55% FRET efficiency in smFRET.

The low spectral dispersion allows us to use sub-pixel localization algorithms, a field that is advancing rapidly^42^, and all future developments are directly applicable to our sSMLM implementation, potentially benefiting from custom deep-learning routines addressing both spectral orders^62,63^. This could open up avenues for better spatial and spectral precision, more information obtained from the 1^st^ order pattern shape, and sSMLM at even higher emitter density. We additionally note that our implementation can readily be combined with SMLM techniques that modulate excitation patterns to increase localization precision^64–69^. Finally, low-dispersion sSMLM as presented here can be combined with three-dimensional SMLM by engineering the PSF in the emission path^70–72^.

Work by Song et al employed sub-pixel localization algorithms for sSMLM, based on data obtained via the – 1^st^ and +1^st^ orders of a non-blazed transmission grating^38^. While this solution is elegant and uses all photons that arrive on the camera chip for both spatial and spectral localization, blocking out the 0^th^ order leads to a significant loss of photons. Moreover, the implementation requires additional optical components (mirrors and lenses) to direct only the −1^st^ and +1^st^ orders on the camera chip. We additionally note that minimizing the spectral dispersion maximises spectral precision by minimization of effects by shot-, background-, and read-out noise^47^.

Further minimization of spectral dispersion in our implementation is possible by using a lower dispersion blazed grating, by decreasing the effective distance of the grating to the camera chip, if the camera housing permits, or by placing the grating close to an intermediate image plane. Whilst this could result in better spectral resolution and higher achievable sSMLM density, a trade-off of this minimization is decreased information content about the shape of the emission spectrum and the risk of imaging the grating itself on the camera. In this study, we used the shape of the emission spectrum only to discriminate FRET states from the donor-only state.

Taken together, we believe that our implementation of low-dispersion sSMLM will find widespread use due to its inherent simplicity and photon efficiency providing access to maximised spatiotemporal and spectral resolution. We further envision applications in which the photon efficient separation of spectrally different entities is desired such as in low-signal flow cytometry. Here, the ideas taken from superresolution microscopy such as sub-pixel localisation and spectral peak determination can be equally applied even for low magnification configurations.

## Acknowledgements

The authors thank current and previous members of our labs for stimulating discussions and ongoing support. We thank Dr. Mattia Fontana for help on the smFRET samples. This manuscript is part of several research projects (#KIEM.K20.01.054 of the research programme NWO KIEM 2020, #18854 of the research programme NWO Take-off phase I), which are (partly) financed by the Dutch Research Council (NWO). K.J.A.M. is funded by a VLAG PhD-fellowship grant awarded to J.H. RR is funded by the European Research Council/Horizon 2020 (ERC-StG-757397) and EA is funded by the MSCA ITN project THERACAT (765497), both awarded to LA. We further acknowledge support from a Road to Innovation grant from the Value Creation Office at Wageningen University & Research.

## Author contributions

Conceptualisation: KM, JH. Data curation: KM. Formal analysis: KM, MG. Funding acquisition: JH, LA. Investigation: KM, MG, NZ, EA, RRB, LA. Methodology: KM, MG, EA, RRB. Project administration: JH. Software: KM, MG. Supervision: NZ, JH. Visualisation: KM. Writing – original draft: KM. Writing – review & editing: All authors.

## Code and data availability

The code used in this manuscript can be found on Github, https://github.com/HohlbeinLab/sSMLMAnalyzer. Data underlying the manuscript is available at http://doi.org/10.5281/zenodo.6778964.

