## Supplementary Information for "Enabling spectrally resolved single-molecule localization microscopy at high emitter densities"

belonging to

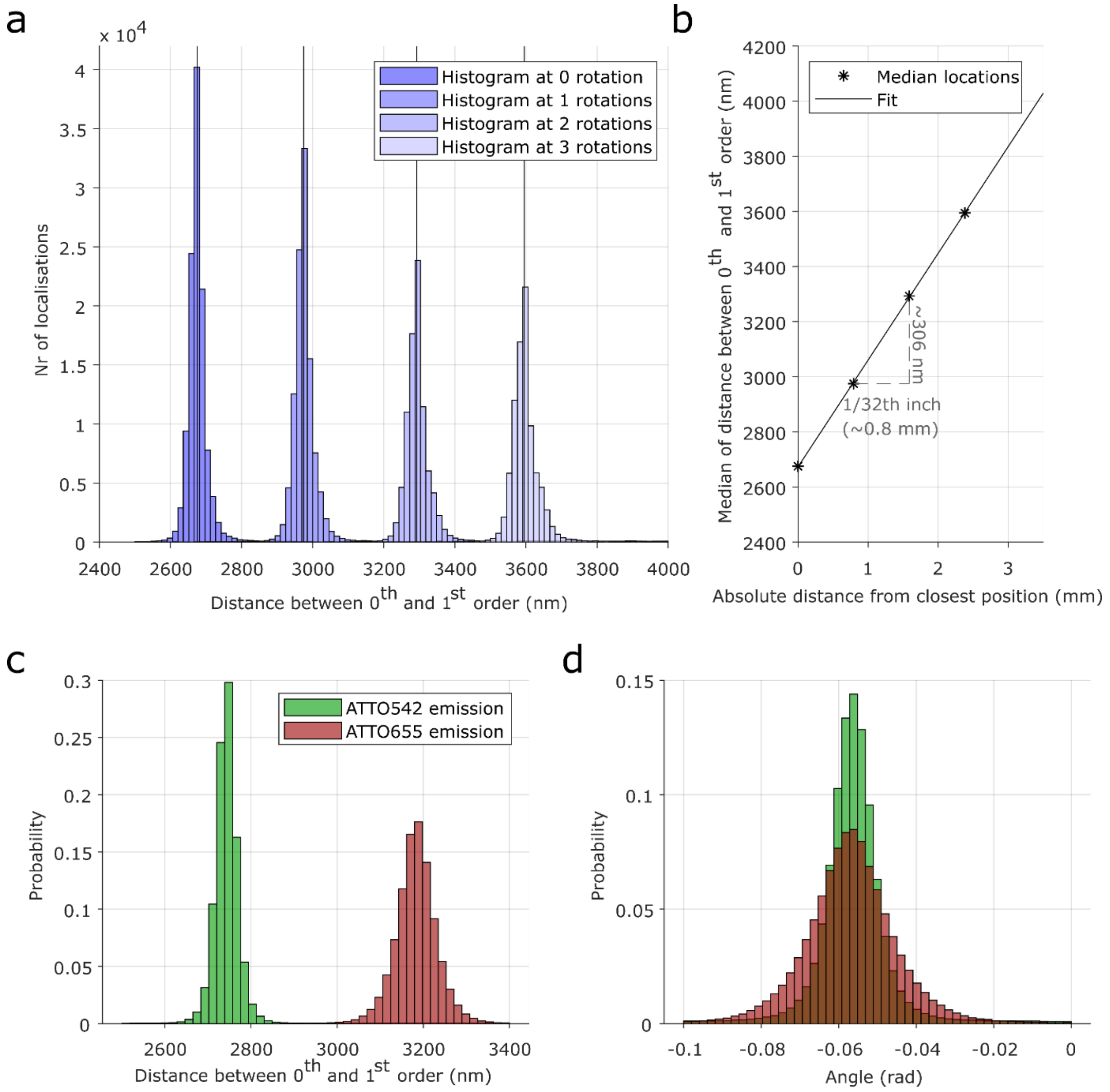


***Supplementary Figure 1***: *Calibration of low-dispersion sSMLM.* ***a,b)*** *Determination of distance between grating and camera chip via the use of a GATTA-PAINT 80RG DNA-PAINT nanoruler (Methods). The grating is incrementally distanced from the camera by a series of rotations (every rotation is 1/32$^{th}$ inch or ~0.8 mm as specified in the utilized c-mount thread). The histograms of the obtained distances between the 0$^{th}$ and 1$^{st}$ order are plotted in* ***a****, while the linear fit of the median distances is shown in shown in* ***b. c,d)*** *Determination of the spectral distance (SD). A DNA-PAINT sample with ATTO542 and ATTO655 fluorophores was imaged. The distances (****c****) show a clear difference between the two fluorophores, while the angle (****d****) is not influenced.*

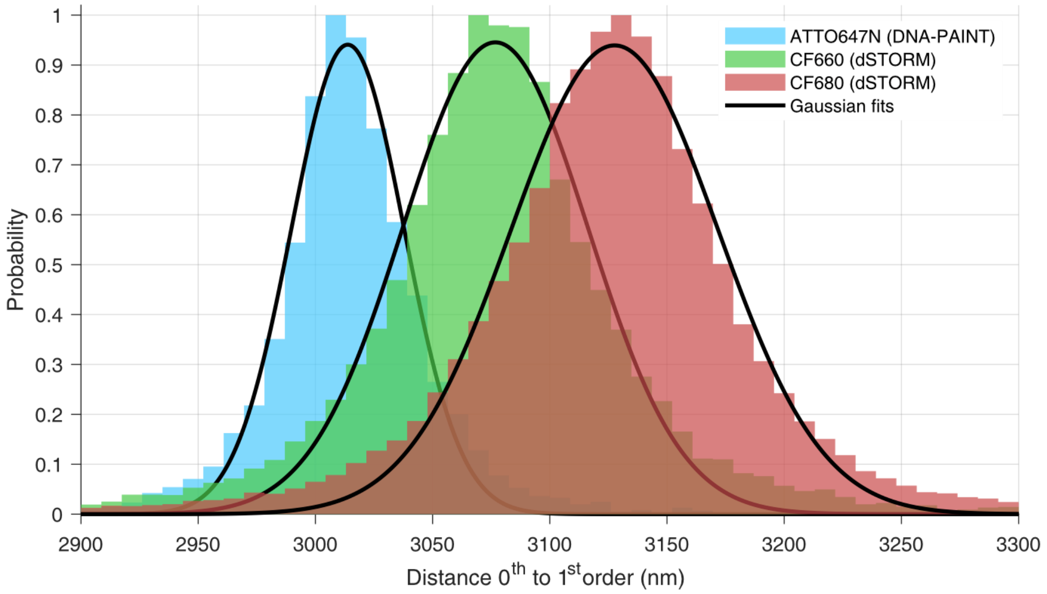


***Supplementary Figure 2****: Technical showcase of triple-fluorophore multiplexing with single-wavelength excitation. Combination of the ATTO647N, CF660, CF680 0th-to-1st-order distance data shown in Figures 2 and 3. Rescaled to provide equal probability, and recoloured for clarity.*

### Supplementary Table 1: ssDNA sequences used for DNA-PAINT on nanoparticles

| Name | Modification | Sequence (5' → 3') |
|---|---|---|
| Imager 1 | ATTO655 (3' side) | CTA GAT GTA T |
| Docking 1 | Biotin (5' side) | TTA TAC ATC TA |
| Imager 2 | ATTO647N (3' side) | TAT GTA GAT C |
| Docking 2 | Biotin (5' side) | TTA TCT ACA TA |

### Supplementary Note 1: Algorithmic determination of pairing distance and rotation

The sSMLM pair finding (explained in more detail in the Methods section) requires user-defined limits for the rotation (orientation of the grading with respect to the camera) and for the expected distances between the PSFs. Here, we discuss a JAVA-based implementation to algorithmically determine the average rotation and distance, and determine boundaries required for the pair finding from there. We note that we cannot determine the distance between the grating and the sensor via this approach, and thus all values are effectively the distance between paired PSFs as measured on the sensor.

Briefly, we run a first FFT (Fast Fourier Transformation) on the reconstructed sSMLM images after single molecule localisation (Supplementary Figure. 3a). The first FFT will produce an image as seen in Supplementary Figure 3b. Cropping the image around the center will lead to a figure that shows a near vertical, strip like pattern representing the periodicity of localisations due to the grating (Supplementary Figure 3c). Next, we apply the directionality functionality of ImageJ to obtain the dominant angle in the image representing the orientation of the grating with respect to the sensor. We then apply this angle inversely to rotate the original image, which leads to all pairs of localisations in the $0^{th}$ and $1^{st}$ order being horizontally aligned.

Starting from the rotated original image, we perform two consecutive FFTs. The first one will return the stripe-like pattern, now oriented vertically (not shown). The second one will determine the major frequencies representing the two expected distanced for our case of having two different fluorophores in our sample (Supplementary Figure 3d). After cropping (Supplementary Figure 3e) and applying some thresholding (thereby keeping the most intense 0.14% of pixels), we can use ImageJ's particle detection feature to determine the position of the peaks. In Supplementary Figure 3f and g, the main features (#5, #2) to the left of the centre peak (#1) represent the two expected distances between the PSFs of the $0^{th}$ and the $1^{st}$ order for the two spectrally distinct fluorophores. The obtained distances will provide the upper and lower estimates for the expected distances then used to run the pair-finding. Here, a secondary angle check is built in as, due to the rotation earlier, the angle of the features to the horizontal should be 0. If this is not the case, a manual check could be necessary. All values are reported to the end user which can then tweak them to process the data using wider or narrower angles/distances.

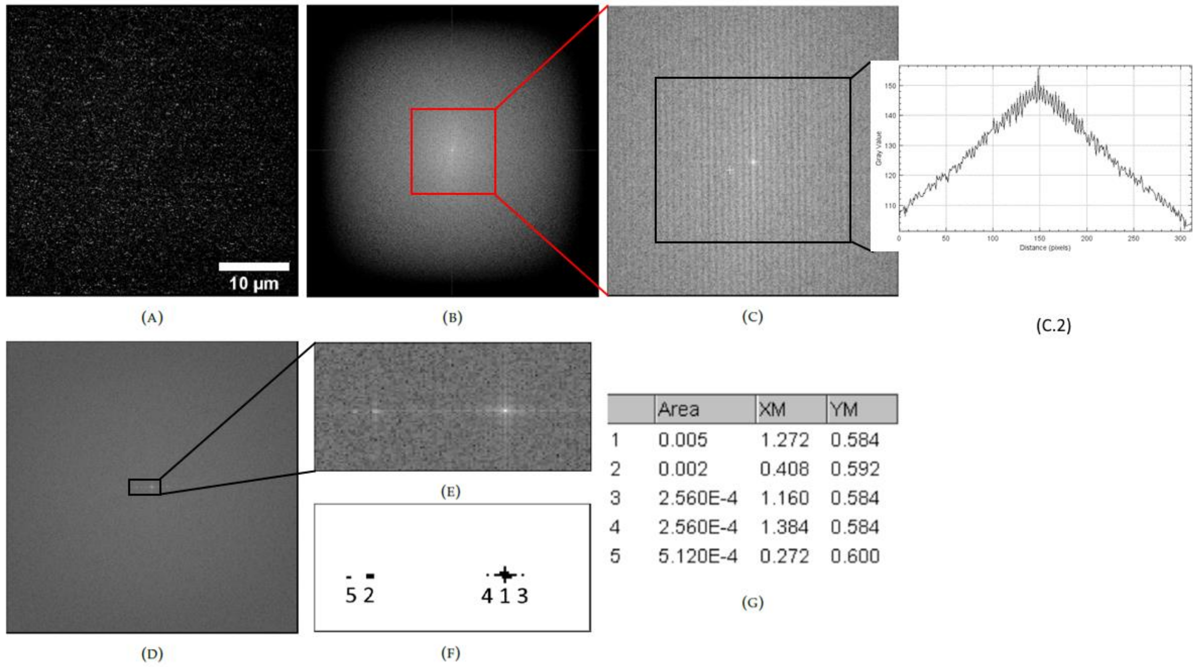


***Supplementary Figure 3:*** *Determining the boundaries for rotation and distances between diffraction orders using Fast Fourier Transformation (FFT).* ***a*** *Super-resolved image of DNA nanorulers (Gattaquant, Germany) with two different fluorophores on the imager DNA strands after analyzing raw data featuring both $0^{th}$ and $1^{st}$ order PSFs with ThunderSTORM-phasor.* ***b*** *2D FFT of* ***a*** *indicates periodicity due to the pairing of single emitters in the 0th and $+1^{st}$ diffraction order.* ***c*** *Zoom-in of* ***b****. From this cropped image the angle is determined using the Directionality functionality in ImageJ. An insert shows the summed plot profile over the marked area, indicating periodicity.* ***b*** *2D FFT of* ***c****. The outer edges of* ***b*** *are discarded as they provide no information in the higher frequency k-space and we want to increase the relative magnitude of the pattern.* ***e*** *Zoom-in of* ***d****, showing the only features of high intensity at the centre. The spots correspond to the wavelengths of the pattern shown in C.2* ***f*** *The same data as in* ***e*** *after thresholding showing only sections of high intensity (top 0.14%).* ***g*** *Running the function 'Analyze Particles' in ImageJ/Fiji on F) determines the position of each particle. The distance from the centre peak (#1) to the features (#2 and# 5) determines the distance boundaries. This also allows for a calculation to check if the angle correction was done properly, since the angle from the features to the centre should be 0 compared to the horizontal.*

The algorithm is furthermore described in pseudocode below.

1. Take the square image containing the sup-pixel localised data (e.g., provided as 2D histogram) and rescale to 1024 pixels by 1024 pixels. Normalise the intensities.
2. A forward FFT is performed using ij.plugin.FFT (size of the resulting image is 1024x1024). After every forward FFT we use ImageJFunctions.wrapNumeric to obtain an image in Cartesian coordinates.
3. Using fiji.analyze.directionality.Directionality_with a binrange of 0 to 180 we obtain the dominant angle, as well as the standard deviation.
4. We rotate the FFT image obtained in 2) by the obtained angle. This is done on the ImageProcessor component. This causes some loss of data since we still maintain a square viewport, but these edges are cropped away later.
5. If the standard deviation of the angle is high (>0.2 rad) we set it to 0.2 rad. Conversely, if it is very low (<0.04 rad) we set it to 0.04 rad. The standard deviation is used to find the angle boundaries (see step 6), so limiting their value ensures that we do not envelop too many points or too few points.

6. We set the upper and lower boundaries for the angle: angle - 2.5 * std to angle + 2.5 * std, respectively. This 2.5 value is set empirically.
7. We crop the rotated FFT image around the center from (256, 256) to (767, 767).
8. On this cropped image another forward FFT is performed. The resolution of the resulting image is 512x512. This image is then cropped from (0, 232) to (256, 281).
9. Using the ImageJ Histogram feature, we determine the value of the top 99.86% of pixels (value determined empirically) and set that as a threshold, In a FFT image, a large amount of pixels are usually of low value (due to low amplitude high frequency components, so this only keeps the few higher intensity features). These isolated features represent the $0^{th}$ order as well as all $1^{st}$ orders of emitters present.
10. On this thresholded image the ImageJ ParticleAnalyser is run, calculating the centers of mass of all features present, as well as the size of the features.
    a. The values in the list are rescaled to the original dimensions
    b. The distance from the feature to the centre of the image, as well as the angle compared to the horizontal '0' are calculated.
    c. Features that have a distance of less than 1% of the width of the image or over 40% of the width are discarded. This limit is set empirically to discard features stretching the whole image or artefacts very close to the centre of the image.
    d. If the angles calculated are over 0.1 rad the angle was not calculated before and is corrected by the angle detected here.
11. If features are found the closest and furthest distances are selected.
    a. The low distance cut-off is the closest point's distance * 0.9 + 0.375% of the width of the image
    b. The high distance cut-off is the furthest point's distance * 1.1 + 0.375% of the width of the image.
12. These values are then used to filter point combinations for each frame.

Many of the values seen have been chosen empirically and have worked well for our applications. These values can be adjusted in the source-code or a custom version of this algorithm can be implemented to then be passed to the plugin in ImageJ's macro feature.
